# The Correlation Between Cell and Nucleus Size is Explained by an Eukaryotic Cell Growth Model

**DOI:** 10.1101/2021.09.01.458491

**Authors:** Yufei Wu, Paul Janmey, Sean X. Sun

## Abstract

In eukaryotes, the cell volume is observed to be strongly correlated with the nuclear volume. The slope of this correlation depends on the cell type, growth condition, and the physical environment of the cell. We develop a computational model of cell growth and proteome increase, incorporating the kinetics of amino acid import, protein/ribosome synthesis and degradation, and active transport of proteins between the cytoplasm and the nucleoplasm. We also include a simple model of ribosome biogenesis and assembly. Results show that the cell volume is tightly correlated with the nuclear volume, and the cytoplasm-nucleoplasm transport rates strongly influences the cell growth rate as well as the cytoplasm/nucleoplasm ratio. Ribosome assembly and the ratio of ribosomal proteins to mature ribosomes also influence the cell volume and the cell growth rate. We find that in order to regulate the cell growth rate and the cytoplasm/nucleoplasm ratio, the cell must optimally control groups of kinetic parameters together, which could explain the quantitative roles of canonical growth pathways. Finally, using an extension of our model and single cell RNAseq data, it is possible to construct a detailed proteome distribution, provided that a quantitative cell division mechanism is known.

**Author summary:** We develop computational model of cell proteome increase and cell growth to compute the cell volume to nuclear volume ratio. The model incorporates essential kinetics of protein and ribosome synthesis/degradation, and their transport across the nuclear envelope. The model also incorporates ribosome biogenesis and assembly. The model identifies the most important parameters in determining the cell growth rate and the nucleoplasm/cytoplasm ratio, and provides a computational starting point to construct the cell proteome based on the RNAseq data.

## Introduction

A universal feature of eukaryotic cells is that the nuclear size appears to be correlated with the cell size [1–3]. This correlation has been quantified in yeast and mammalian cells, cells in epithelial tissues [4], and cells growing in different nutrient and physical environments (Fig. 1(a)). It occurs in a population of isogenic cells as well as across populations of different cell types. This correlation also persists in normal as well as cancer cells, although oncogene mutations appear to alter the nucleoplasm-cytoplasm ratio. In growing cells, the cytoplasm is continuously expanding from amino-acid and nutrient import, and also protein synthesis, but is unclear why the nucleoplasm expands proportionally and whether the constant nuclear-cell size ratio serves an important functional role. In this paper, we explore the question of nucleus-cell size correlation using a phenomenological mathematical model. We find that by incorporating a model of ribosome biogenesis, we can quantitatively explain cell-nuclear size correlation and the N/C ratio. Moreover, our model suggests that nucleoplasm-cytoplasm transport plays an important role in the overall growth of the cell. A balanced proteome is necessary to achieve optimal growth control.

**Fig 1.**
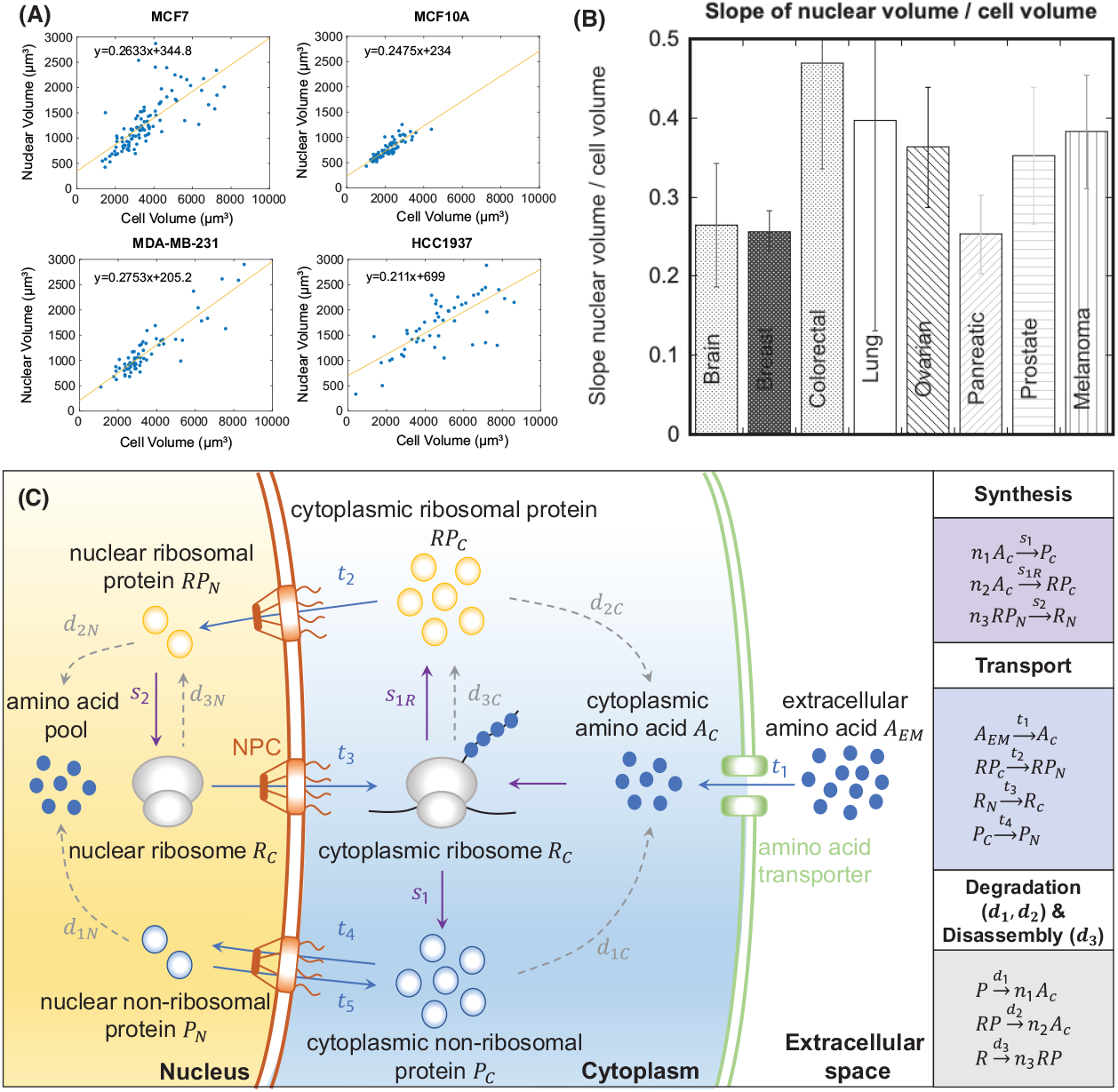
(A) Nuclear volume vs. cell volume for several cell lines grown on 500Pa polyacrylamide (PAA) substrates coated with collagen I. Each point is a single cell. The nuclear volume is proportional to the cell volume. (B) Slope of nuclear volume vs. cell volume for cell lines from several different types of human tissue. Brain (T98G, U-87), breast (MCF10A0JSB, HTERT-HME1, MDA-MB-231, MCF7, HCC1937a, T-47D), colorectal (SW480, SW620, HT-29, HCT116), lung (NCI-H2087, NL20, NCI-H2126), ovary (SK-OV-3, NIH: OVCAR-3, Coav3), pancreas (Panc-1, HTERT-HPNE, Capan-1), prostate (DU145, 22RV1, LnCaP Clone FG, PC-3, RWPE-1), skin (WM266-4, A375, MeWo, SK-MEL-2). (C) A model of eukaryotic cell growth with mass transport. During cell growth, amino acids are imported into the cytoplasm and assembled into non-ribosomal proteins and ribosomal proteins by mature ribosome particles. The ribosomal proteins are transported into the nucleoplasm to combine with ribosomal RNA to mature into ribosome particles, which are then transported back out to the cytoplasm. Non-ribosomal proteins are also actively transported in and out of the nucleoplasm. Proteins and ribosomes are also actively degraded. These processes can be captured by a simple set of mass flux equations.

For any cell, the overall cell mass increase must come from the net import of extracellular material: water, amino-acids, nutrients, endocytosed and phagocytosed materials, etc. In the cytoplasm, mature ribosome complexes are responsible for protein translation and generation of new proteins. For eukaryotes, ribosomes undergo a maturation process [5, 6] where new ribosomal proteins are made in the cytoplasm, and then are transported into the nucleoplasm to combine with ribosomal RNA to form large and small subunits. Once matured, the ribosomal subunits are transported out of the nucleus into the cytoplasm and combine into a complete ribosome to make new proteins from mRNA. The transport process across the nuclear envelope is carried out by the RanGTPase cycle, which also transports other proteins across the nuclear envelope [7]. Therefore, the cytoplasmic population of mature ribosomes depends on nuclear-cytoplasmic transport, which in turn determines the protein synthesis rate.

Using a simple model, we demonstrate that the rate of nuclear-cytoplasmic transport directly influences the nuclear-cytoplasm ratio, and the cell growth rate. The model is not molecularly detailed, i.e., it does not model individual proteins and their interactions. Rather it models the overall population of proteins and ribosomes, and therefore is a coarse-grained cell scale model. Refinement of the model framework and connections to gene regulation models are discussed.

We begin by outlining equations governing the time evolution of the cell proteome number together with ribosome production. We then consider the role of the nucleoplasm-cytoplasm transport in ribosome biogenesis and incorporate this element in the minimal model. We also consider proteolysis or protein degradation. The solution of these equations predicts an exponential increase in assembled proteins, ribosomal proteins, and mature ribosome particle. Moreover, the nucleoplasm and cytoplasm components of proteins and ribosomes all increase at the same rate, and therefore the model predicts that the nuclear volume is proportional to the cytoplasmic volume. We analyze several regimes of the model, including when the proportion of the mature ribosome and ribosomal proteins is altered, such as during aneuploidy. The results show that the right proportion of mature ribosome particles is needed to maintain a high cell growth rate. By analyzing the gradient of growth rate and C/N ratio with respect to model parameters, it is found that to control cell growth rates, it is necessary to not only change amino-acid import and protein synthesis rates, but also modulate nucleoplasm-cytoplasm transport.

### A Cell Growth Model With Ribosome Biogenesis and Nucleoplasm-Cytoplasm Transport

A schematic of the model is shown in Fig. 1(b). In this model, we consider essential processes in an eukaryotic cell, which includes protein synthesis, transport across various compartments, protein degradation and ribosome disassembly. For simplicity, we assume that all the molecules and particles are evenly distributed in cytoplasm and nucleus, so we can use a constant concentration to describe each component. In addition, we assume that the cell volume is proportional to the number of macromolecules (including proteins and ribosomes), which is a good approximation for mammalian cells [8]. Under these assumptions, we can treat all the metabolic activities as chemical reactions in an isobaric and isothermal compartment. All the variables considered contain subscripts, which indicates the cytoplasmic (*C*) and nucleoplasm (*N*) compartments. 1) *P*_*C,N*_ are the number of non-ribosomal proteins in the cytoplasm and nucleoplasm, respectively. 2) *A*_*C*_, the number of amino acid molecules in the cytoplasm. *RP*_*C,N*_ are the number of ribosomal proteins in the cytoplasm/nucleoplasm. *R*_*C,N*_ are the assembled mature ribosome particles in the cytoplasm/nucleoplasm. The model variables and their order-of-magnitude estimates are given in Table I.

**Table 1.** Glossary of model variables and their numerical estimates for new born cells. See text for references.

| Symbol | Description | Yeast Values | Mammalian Values | Source |
| --- | --- | --- | --- | --- |
| $A_C$ | Cytoplasmic amino acid (AA) number | $3200 \times 10^6$ | $21600 \times 10^6$ | [16, 29, 50, 51] |
| $P_C$ | Cytoplasmic non-ribosomal protein number | $15 \times 10^6$ | $660 \times 10^6$ | [25, 30, 31, 52] |
| $R_C$ | Cytoplasmic mature ribosome number | $0.135 \times 10^6$ | $6 \times 10^6$ | [25, 53] |
| $RP_C$ | Cytoplasmic ribosomal protein number | $1.35 \times 10^6$ | $60 \times 10^6$ | [25, 30, 31, 52] |
| $RP_N$ | Nuclear ribosomal protein number | $0.15 \times 10^6$ | $15 \times 10^6$ | [25, 30, 31, 52] |
| $R_N$ | Nuclear ribosome complex number | $0.015 \times 10^6$ | $1.5 \times 10^6$ | [25, 53] |
| $P_N$ | Nuclear non-ribosomal protein number | $1.65 \times 10^6$ | $165 \times 10^6$ | [25, 30, 31, 52] |
| $V_C$ | Cytoplasmic volume ($\mu\text{m}^3$ or fL) | 27 | 1500 | [26, 28, 55, 56] |
| $V_N$ | Nuclear volume ($\mu\text{m}^3$ or fL) | 3 | 500 | [26–28, 54–56] |

### Model Component 1: Protein Synthesis

Protein synthesis is a sequential assembly process in the cytoplasm by mature ribosome particles. At each step, a new amino acid is added to a peptide chain (which is attached to the ribosome) and then the ribosome moves to the next codon (Fig. 2). We assume that each step takes the same length of time and the total steps (average amino acid number) for non-ribosomal protein synthesis and ribosomal protein synthesis are *n*_1_ and *n*_2_, respectively. In a sequential assembly process, the synthesis rate is determined by the last step where *n*_*i*_ − 1 peptides become *n*_*i*_ (*i* = 1, 2) peptides [9]. The synthesis rate is proportional to concentration of peptide chains with *n*_*i*_ − 1 (*i* = 1, 2) amino acids. If we assume there is sufficient tRNA, the synthesis rate should also be proportional to amino acid concentration. Accordingly, we can write the rate of change in non-ribosomal protein number in terms of their synthesis rate in a container with volume *V*_*C*_ as [10]:

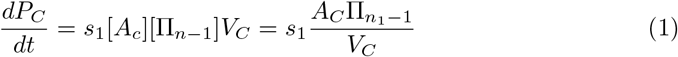

where *P*_*C*_ is cytoplasmic non-ribosomal protein number, *A*_*C*_ is cytoplasmic amino acid number, 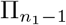 is the number of peptide chains in the last synthesis step, *V*_*C*_ is the cytoplasm volume and *s*_1_ is the synthesis rate coefficient (which depends on the protein mRNA, the ribosome translation speed and biochemical signaling networks that control synthesis). The brackets indicate their molar concentrations. 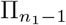 can be estimated as follows. For *R*_*C*_, the number of mature ribosomes, some portion of which are carrying out protein synthesis, assuming that peptide chain lengths in the ribosomes during active synthesis are randomly distributed, the probability of finding peptide chains at the last step is 1*/n*_1_. Therefore on average, 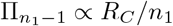. The protein synthesis rate can be expressed as:

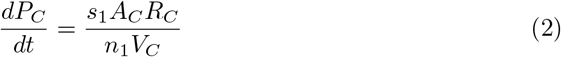

**Fig 2.**
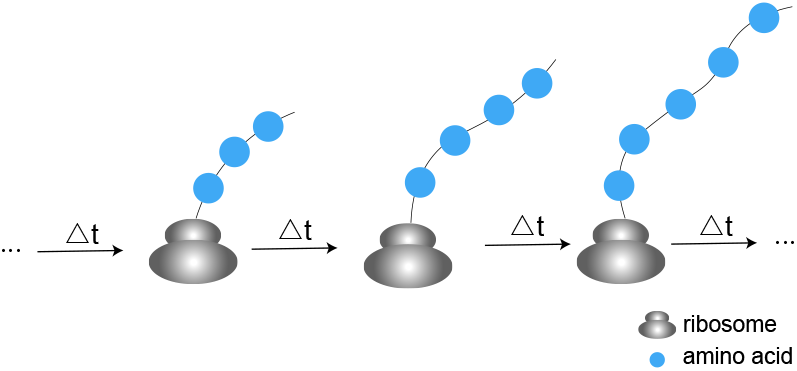
Protein synthesis is a sequential assembly process carried out by mature ribosomes. The rate of protein synthesis is proportional to the concentrations of amino acid and ribosome-peptide complex at the last assembly step (see Eq. (2)).

Similarly, we can write the synthesis equation for ribosomal protein *RP*_*C*_ (with synthesis rate coefficient *s*_1*R*_ and average amino acid number *n*_2_ in a protein).

For ribosome assembly and maturation in the nucleus (*R*_*N*_), we can assume there are several maturation sites (nucleolus), whose number and size are proportional to the nuclear volume (*V*_*N*_), and the assembly should also be sequential. Therefore, similar to protein synthesis, we can write the ribosome assembly equation as:

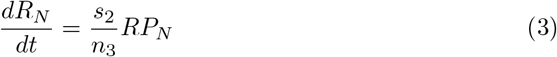

where *RP*_*N*_ is the ribosomal protein number in the nucleus, *n*_3_ is the average ribosomal protein number in a ribosome and *s*_2_ is the synthesis coefficient, including the number of synthesis sites and the assembly rate. According to the stoichiometric coefficients (Fig. 1(b)), the rate of amino acid (*A*_*C*_) and nuclear ribosomal protein (*RP*_*N*_) loss due to synthesis can be written as:

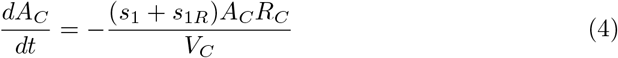

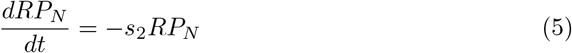

### Model Component 2: Transport

To synthesize proteins, the cell needs to import amino acids from the extracellular environment, and some of the imported amino acids go into the nucleus via free diffusion. After completing synthesis, cytoplasmic ribosomal proteins are transported into the nucleus to assemble into ribosomal subunits. Then the two assembled ribosomal subunits are transported out of the nucleus, combine into a mature ribosome and participate in protein translation [5, 6]. For simplicity, in this model, we assume the complete ribosome is directly assembled in the cell nucleus. There is also bidirectional non-ribosomal protein transport across the nuclear envelope. Most of the macromolecule transport fluxes in the cell are active. For amino-acids in the cytoplasm, amino acid transporters (Slc gene family) on the cell surface are responsible for their import [11]. For protein transport across the nuclear envelope, import and export are carried out by the RanGTPase cycle [7].

In general, the transport rate depends on both the cargo concentration and transport protein density. For amino-acid import in our model, we will consider poor and rich nutrient conditions, corresponding to transporter saturated and cargo saturated conditions, respectively. For protein and ribosome transport across the nuclear envelope, we assume both the cargo and transporter proteins are unsaturated. If we assume that the transporter protein number (*TP*_*C*_) is proportional to total protein number in cytoplasm *P*_*C*_, we can obtain a general equation for active transport of protein and ribosome (when cargo concentration and transport protein density are both unsaturated):

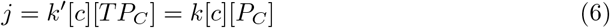

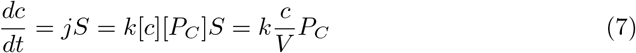

Where *j* is the cargo flux through membrane (per unit area per unit time), [*c*], [*TP*_*C*_] and [*P*_*C*_] are concentration of cargo, transporter protein and cytoplasmic protein, respectively. *c* is the total cargo number, *S* is the surface area. The number of transport proteins is the concentration times the area of the nuclear envelope, which is proportional to *P_C_*, i.e., 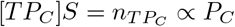. *k* is the transport rate, which also includes the percentage of transport proteins. Based on this general equation, we can write all the transport equations as:

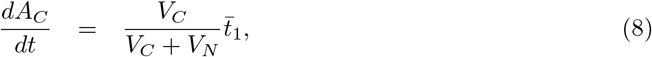

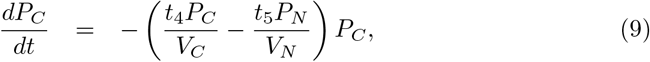

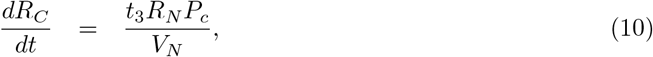

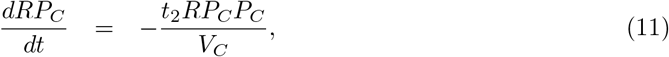

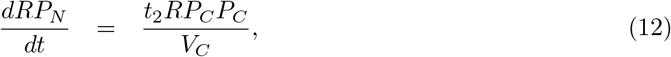

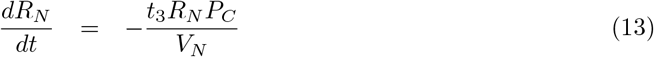

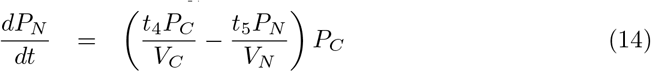

*P*_*N*_ is the nuclear non-ribosomal protein number, *t*_2_ is the import coefficient of cytoplasmic ribosomal protein, *t*_3_ is the export coefficient of nuclear ribosome, *t*_4_ and *t*_5_ are import and export coefficients of non-ribosomal protein through the nucleus. 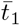 includes the exterior amino acid concentration and the import coefficient of amino acid. 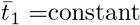 when exterior amino acids are poor while 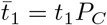 when exterior amino acids are rich. For simplicity, we neglect the process of amino acid diffusion across nuclear envelop and assume that the imported amino acids are evenly distributed in cytoplasm and nucleoplasm depending on *V*_*C*_*/V*_*N*_.

### Model Component 3: Degradation and disassembly

In this part, we consider protein degradation and ribosome disassembly, which occurs in both nucleus and cytoplasm [12]. The degraded proteins are broken down into short peptide chains and amino acids, and they are recycled in new protein translation.

Similarly, ribosome disassembly produces new ribosomal proteins, which can also be reused in the ribosome synthesis cycle (Fig. 1(b)). For simplicity, we assume that all the proteins only break down into amino acids, and these amino acids are evenly distributed in cytoplasm and nucleoplasm depending on the C/N ratio. There are two major pathways of protein degradation: the ubiquitin-proteasome pathway and the lysosomal proteolysis-mediate protein degradation [12, 13]. In both pathways, degradation is aided by degradation proteins or protein complexes. Therefore, the degradation rate is proportional to concentrations of degradation proteins and proteins to be degraded. Assuming the numbers of proteins assisting degradation in cytoplasm and nucleus are proportional to the numbers of non-ribosomal proteins in cytoplasm and nucleus respectively, we can get the general expression for protein degradation:

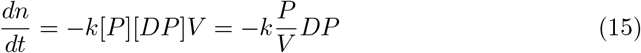

Where [*P*] and [*DP*] are concentrations of proteins to be degraded and assisting degradation. Similarly, we can obtain the disassembly equation for ribosomes.

Accordingly, we can write all the degradation and disassembly equations:

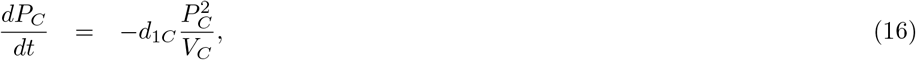

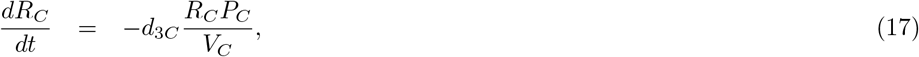

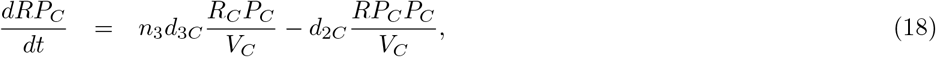

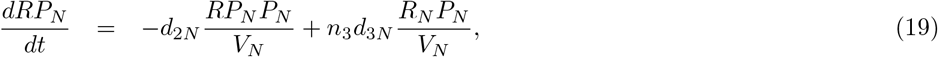

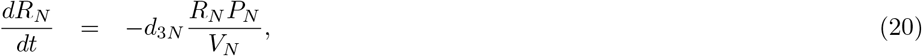

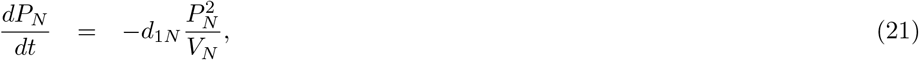

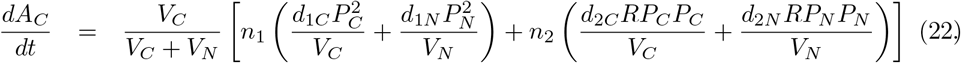

*d*_1*C*_,*d*_1*N*_ are the degradation coefficients of non-ribosomal proteins in cytoplasm and nucleus. *d*_2*C*_,*d*_2*N*_ denote the degradation coefficients of ribosomal proteins. *d*_3*C*_, *d*_3*N*_ are the disassembly coefficients of ribosomes. These coefficients include both the degradation rate and percentage of degradation proteins assisting the process.

Putting all three parts together, we obtain the governing equations for the dynamics of the complete proteome (See parameter details in Table 2):

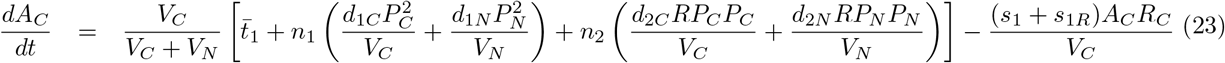

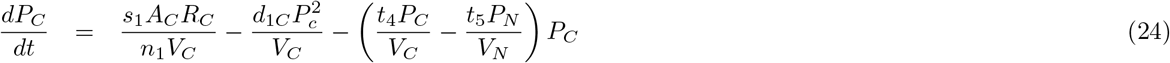

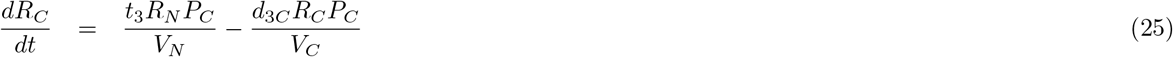

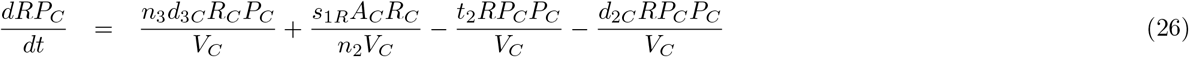

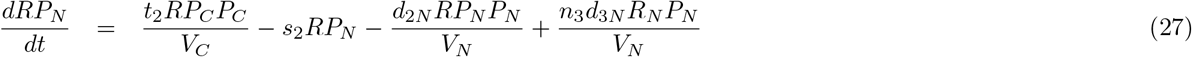

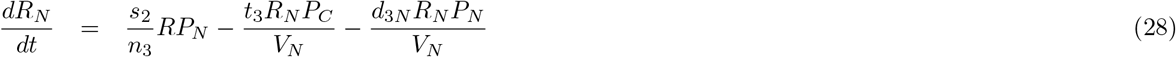

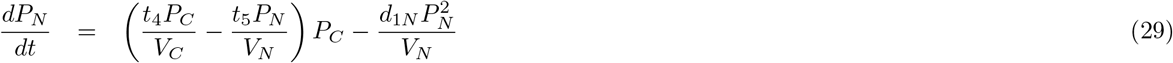

where *A*_*C*_, *P*_*C,N*_, *R*_*C,N*_, *RP*_*C,N*_ are cytoplasmic/neoplasmic number of amino acids, non-ribosomal proteins, ribosomes and ribosomal proteins, respectively. *n*_1_, *n*_2_ are average amino acid numbers per non-ribosomal and ribosomal protein. *n*_3_ is the average number of proteins in a ribosome. 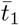 is the amino acid uptake rate, which includes the exterior amino acid abundance, transport efficiency and percentage of transport proteins. 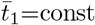 when exterior amino acid is insufficient while 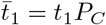 when amino acid is sufficient. *t*_2_, *t*_3_, *t*_4_, *t*_5_ are transport coefficients of *RP*_*C*_, *R*_*N*_, *P*_*C*_, *P*_*N*_, which include transport efficiency and percentage of transport protein. *d*_1*C*_, *d*_2*C*_, *d*_1*N*_, *d*_2*N*_, *d*_3*C*_, *d*_3*N*_ are degradation or disassembly coefficients, which include degradation efficiency and percentage of degradation (assisting) proteins. *s*_1_ and *s*_1*r*_ are synthesis coefficients of non-ribosomal and ribosomal protein, which include mRNA concentration and ribosome moving speed. *s*_2_ is the assembly coefficient of ribosome, which includes the number of assembly sites and assembly rate at each site. *V*_*N*_ and *V*_*C*_ are nucleus and cytoplasm volumes, respectively.

**Table 2.** Model parameters and their numerical estimates.

| Parameter | Description | Yeast | Mammalian | Source |
| --- | --- | --- | --- | --- |
| $r_{1,2}$ | Macromolecule number and cytoplasmic (nuclear) volume ratio ($10^6/\mu\text{m}^3$) | 0.6 | 0.6 | Estimate |
| $n_1$ | Average number of AA in non-ribosomal proteins | 400 | 400 | [32] |
| $n_2$ | Average number of AA in ribosomal proteins | 400 | 400 | [32] |
| $n_3$ | Average number of protein subunits in mature ribosomes | 80 | 80 | [49] |
| $t_1$ | Amino acid import rate coefficient ($\text{h}^{-1}$) | 400 | 36 | Estimate |
| $t_2$ | Ribosomal protein transport coefficient across nuclear envelop ($\mu\text{m}^3/10^6\text{h}$) | 9 | 1 | [33] |
| $t_3$ | Ribosome transport coefficient across nuclear envelop ($\mu\text{m}^3/10^6\text{h}$) | 1.2 | 0.15 | [33] |
| $t_4$ | Non-ribosomal protein transport coefficient into nucleus ($\mu\text{m}^3/10^6\text{h}$) | 0.8 | 0.1 | Estimate |
| $t_5$ | Non-ribosomal protein transport coefficient out of nucleus ($\mu\text{m}^3/10^6\text{h}$) | 0.75 | 0.05 | Estimate |
| $d_{1C}, d_{1N}$ | Non-ribosomal protein degradation coefficient ($\mu\text{m}^3/10^6\text{h}$) | 0.1 | 0.01 | Estimate |
| $d_{2C}, d_{2N}$ | Ribosomal protein degradation coefficient ($\mu\text{m}^3/10^6\text{h}$) | 0.1 | 0.01 | Estimate |
| $d_{3C}, d_{3N}$ | Ribosome disassembly coefficient ($\mu\text{m}^3/10^6\text{h}$) | 0.1 | 0.01 | Estimate |
| $s_1$ | Non-ribosomal protein synthesis coefficient ($\mu\text{m}^3/10^6\text{h}$) | 155 | 115 | [25, 30, 31] |
| $s_{1R}$ | Ribosomal protein synthesis coefficient ($\mu\text{m}^3/10^6\text{h}$) | 125 | 95 | [25, 30, 31] |
| $s_2$ | Ribosome assembly coefficient ($\text{h}^{-1}$) | 48 | 1.5 | [25] |

### On the Mapping Between Molecular Content and Cell and Nucleus Size

Equations 23-29 are the general equations for the time rate of change of protein, ribosomal protein, ribosome particles and amino acids in connected cytoplasm and nucleoplasm compartments with volumes *V*_*C*_ and *V*_*N*_, respectively. The remaining important question is how are *V*_*C*_ and *V*_*N*_ determined. The largest number of molecules in the cell is water, followed by ions [14]. The number of amino acids and proteins are smaller in comparison, although in yeast the amino-acid concentration is comparable to ionic concentration [15, 16]. From studies on cell ion regulation, it is found that by actively passaging ions across the cell surface, cells maintain a constant osmolarity difference between the cytoplasm and the extracellular environment [17–19]. This ion homeostasis model may be generalized to include charged small organic molecules such as some amino-acids, whose import must rely on transporters that are either sensitive to membrane voltage, or co-transporters that import these charged molecules with their associated counter ions (Fig. 3) [11, 20]. Therefore, it is reasonable to consider ions, amino-acids and other charged small molecules as part of the cell ionic homeostasis system.

**Fig 3.**
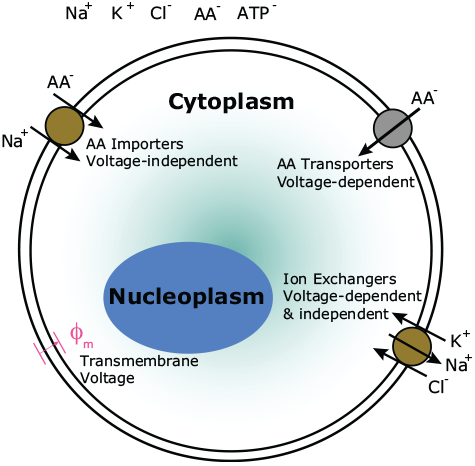
The cell volume is proportional to the impermeable macromolecular number (proteins, buffers, and ribosomes) because the concentration of small charged molecules such as amino acids and ATP are likely regulated together with concentration of ions. For example, amino acids can be imported into the cytoplasm together with Na^+^ in a voltage independent manner, or imported individually in a voltage dependent manner. The transmembrane voltage *ϕ*_*m*_ also depends on ion concentration (Na^+^, K^+^, Cl^−^, etc). Models of cell ionic homeostasis predict that the cell water content is directly proportional to cell macromolecular number.

As a consequence of the cell ion homeostasis, the water content is proportional to the total number of *impermeable* molecules such as proteins and buffers in the cytoplasm [19, 21]. Ions and small amino acids are thought to freely diffuse across the nuclear envelope according to their concentration gradients. Therefore, at equilibrium, there is no ion and small molecule concentration differences between the nucleoplasm and cytoplasm, but there could be a protein concentration difference.

If we accept that the concentrations of small molecules and ions are regulated together, the it is reasonable to assume that the cell volume is proportional to the protein and macromolecular number.

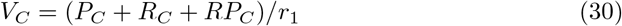

where *r*_1_ is a proportionality constant that may depend on the ionic regulation system and the overall osmotic pressure of the cytoplasm. The total osmotic pressure difference between the cytoplasm and the external environment also should depend on the cytoplasmic hydraulic pressure, because at equilibrium the osmotic pressure difference is equal to the hydraulic pressure difference. The hydraulic pressure, in turn, may depend on cytoskeletal contractility, and therefore *r*_1_ may depend on other variables such as cytoskeletal activity.

For the nucleoplasm, we have

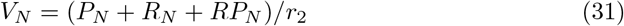

Here we see that because the total osmolarity of the nucleus may be different than the cytoplasm, there might be an osmotic gradient and hydraulic pressure gradient across the nuclear envelope. This excess pressure in the nucleus must be mechanically balanced by the tension in the nuclear envelope and/or cytoskeleton activity. Indeed, for mammalian cells, F-actin and associated contractile forces are linked to the nuclear lamins through the LINC complex [22]. Therefore, there could be mechanical tension in the nuclear envelope. Moreover, intermediate filaments such as vimentin also surrounds the nucleus and may provide additional mechanical support [23]. Therefore, the proportionality constants, *r*_1,2_, between the volume of the compartment and the macromolecular content could be different. The nuclear compartment also contains DNA, which in principal also contributes to volume. However, the overall molecular weight of DNA is negligible when compared to protein and ribosome mass [24] and therefore we neglect it here.

Another complicating feature of the volume model is that the nuclear ribosomes are not well mixed with the nucleoplasm, and instead are phase separated into nucleolus. The contribution of nucleolus to the nuclear volume is unclear, and a different model utilizing the volume of phase separated material might be needed. Such a model may be different in numerical detail, but the overall nuclear volume should still be generally proportional to the volume of phase separated nucleolus.

### Parameter Estimation

All initial parameter estimates are based on *Saccharomyces cerevisiae*. The cell cycle time for yeast is around 2 hours [25]. A typical cell volume is 40 *µ*m^3^ [26], and cell to nuclear volume ratio (C/N ratio) is around 10 [27]. Assuming that mature cells are 60 *µ*m^3^ [28] and newborn cells grow exponentially from half of the mature size: 30 *µ*m^3^, we can obtain a rough estimation for cell volumeJduring a cell cycle: *V*_*C*_ ≈ 30*e*^0.35*t*^*µ*m^3^. We can also obtain an average cell size: 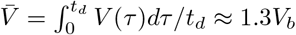, where *V*_*b*_ is the volume of a newborn cell. The following calculations will be based on this average cell state (All components are assumed proportional to cell volume).

For mammalian cells, fewer parameters are available from literature. Here we extrapolate some of the parameters from yeast and scale up.

### Initial conditions

Intracellular concentration of amino acids could vary from 100 to 500mM for yeast cells [16, 29]. In our estimate, we set the concentration to be 200mM, so the total number of amino acids for a newborn cell is: *A*_*C*_ ≈ 3.2 ×10^9^. The ribosome number for a newborn cell is around 0.15 ×10^6^ [25]. Assuming the ribosome numbers in cytoplasm and nucleus are proportional to cytoplasm and nucleus volume, we obtain:

*R*_*C*_ = 0.135 × 10^6^, *R*_*N*_ = 0.015 × 10^6^. A mature yeast cell has a total of around 6 × 10^7^ proteins [30], which contains both ribosomal and non-ribosomal proteins. Therefore, for a new born cell, we have approximately 1/2 of the mature cell:

*n*_3_*R*_*C*_ + *RP*_*C*_ + *P*_*C*_ = 27× 10^6^ for cytoplasmic proteins, and *n*_3_*R*_*N*_ + *RP*_*N*_ + *P*_*N*_ = 3 × 10^6^ for nuclear proteins, where *n*_3_ = 80 is the average number of proteins in a mature ribosome particle. Among all the proteins, ribosomal protein percentage could vary from 30% to 50% [25, 31]. In our estimate, we set the fraction to be 45%. Assuming the amount in the cytoplasm is 9 times of that in the nucleus, we have: *RP*_*C*_ = 0.45 27 × 10^6^ − 80× 0.135 × 10^6^ ≈ 1.35 × 10^6^, *P*_*C*_ = 27 × 10^6^− 80 ×0.135 × 10^6^− 1.35 10^6^ = 1.5× 10^7^, *RP*_*N*_ = 0.15 × 10^6^, *P*_*N*_ = 1.65 ×10^6^.

The conversion factor between macromolecule number and volume therefore, is: (*R*_*C*_ + *RP*_*C*_ + *P*_*C*_)*/V*_*C*_ ≈ 0.6 ×10^6^*/µ*m^3^. We assume for both yeast and mammalian cells, this conversion factor is the same for the nuclear compartment.

### Kinetic Parameters

We first consider parameters in synthesis. We assume that the average numbers of amino acids in each ribosomal protein and non-ribosomal protein are the same (*n*_1_ = *n*_2_), which is around 400 [32]. Roughly speaking, a yeast cell must produce 2000 ribosomes per min [25] to maintain metabolism, which means: *s*_2_*/n*_3_*RP*_*N*_ = *s*_2_*/*80 * 0.15× 10^6^*/*0.75 ≈ 0.12× 10^6^h^*−*1^, so *s*_2_ = 48 h^*−*1^. There is a factor of 0.75 because we are considering the average cell state. Assuming newborn cells have half of the mature cell proteins, the cell needs to synthesize 3 10^7^ proteins in a single cell cycle, thus: ((*s*_1_ + *s*_1*r*_)*A*_*C*_*R*_*C*_)*/*(*n*_1_*V*_*C*_) = 1.5 ×10^7^h^*−*1^, so *s*_1_ + *s*_1*r*_ = 280*µ*m^3^*/*10^6^h. Synthesis rate constant should be positively correlated to the amount of each component, so for simplicity, we assume *s*_1_ : *s*_1*r*_ = *P* : *RP* = 11 : 9. Then we obtain: *s*_1_ = 155, *s*_1*r*_ = 125*µ*m^3^*/*10^6^h.

For parameters in transport, we first consider the amino acid uptake rate in rich-nutrient (sufficient amino acids) case. The imported amino acids are utilized in two parts: synthesizing proteins and maintaining the amino acid pool concentration, so *t*_1_≈*P*_*C*_ = *n*_1_((*s*_1_ + *s*_1*r*_)*A*_*C*_*R*_*C*_)*/*(*n*_1_*V*_*C*_) + (*A*_*C*_(*t*) − *A*_*C*_(0))*/*2h= 6000 + 1800. Therefore: *t*_1_ ≈ 400h^*−*1^. During the ribosome assembly process, a rapidly growing yeast cell must import ∼150000 ribosomal proteins per minute across the nuclear envelope, and export ∼4000 ribosomal subunits (equivalently 2000 mature ribosomes) per minute [33], which means: *t*_2_*RP*_*C*_*P*_*C*_*/V*_*C*_ = 9 ×10^6^*/*h, (*t*_3_*R*_*N*_ *P*_*C*_)*/V*_*N*_ = 0.12 ×10^6^*/*h, so *t*_2_ = 9*µ*m^3^*/*10^6^h, *t*_3_ = 1.2*µ*m^3^*/*10^6^h. For non-ribosomal protein transport through the nuclear envelope, we assume that the transport rate is equal to that of ribosomal protein: 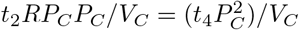, so *t*_4_ = 0.8*µ*m^3^*/*10^6^h. The net import should be positive, so we obtain *t*_5_ = 0.72, which is slightly lower than *t*_4_.

For protein degradation and ribosome disassembly, we assume ribosome disassembly and protein degradation rates in rapidly growing cells are 1*/*10 of the synthesis rate: 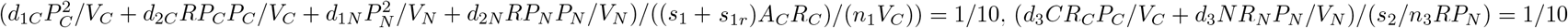.

Assuming all the degradation rates are equal, we can get: *d*_1*C*_ = *d*_2*C*_ = *d*_1*N*_ = *d*_2*N*_ = 0.1*µ*m^3^*/*10^6^h, *d*_3*C*_ = *d*_3*N*_ = 0.1*µ*m^3^*/*10^6^h. A summary of all estimated kinetic parameters is shown in Table 2.

For mammalian cells, the cell volume is roughly 100 times larger, and the C/N ratio is around 3-10. Typical cell cycle time is around 20 hours. Therefore, the parameters are different. Our estimates for the parameters for a prototypical mammalian cell are also given in Table 2.

## Results

### Exponential vs linear growth are determined by amino acid import

Using the proposed models of *V*_*C*_ and *V*_*N*_ in Eqs. (30, 31) and the estimated parameters, we can compute the change in cell protein content, ribosome content, nuclear volume, and the cell volume during a complete cell cycle for both yeast and mammalian cells in different conditions (e.g. rich amino acid, poor amino acid, and quiescent cases). In the simulation, the cell divides when the cell volume reaches twice the initial cell volume *V*_0_. After division, all the components are set to 1*/*2 of the mother cell number. We note that for *S. cerevisiae*, the mother cell grows a bud at replication initiation, and the volume of the bud after pinching off from the mother cell can be variable. At best nutrient conditions, the daughter cell volume can reach approximately 1/2 of the mother cell [34]. The division mechanism our simulation does not effect cell growth rate or the computed cytoplasm to nucleoplasm ratio. In the case where the amino acid import rate is proportional to the protein number, i.e., 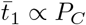, our model predicts exponential growth of cell components in terms of their overall number, and exponential growth of cell and nuclear volume (Fig. 4(a,b,e,f)), i.e., all quantities are proportional to *e*^*λt*^, where *λ* is the cell growth rate. For these parameters, the majority of the cell consists of non-ribosomal proteins, although ribosomal proteins is a substantial fraction of the overall proteome.

**Fig 4.**
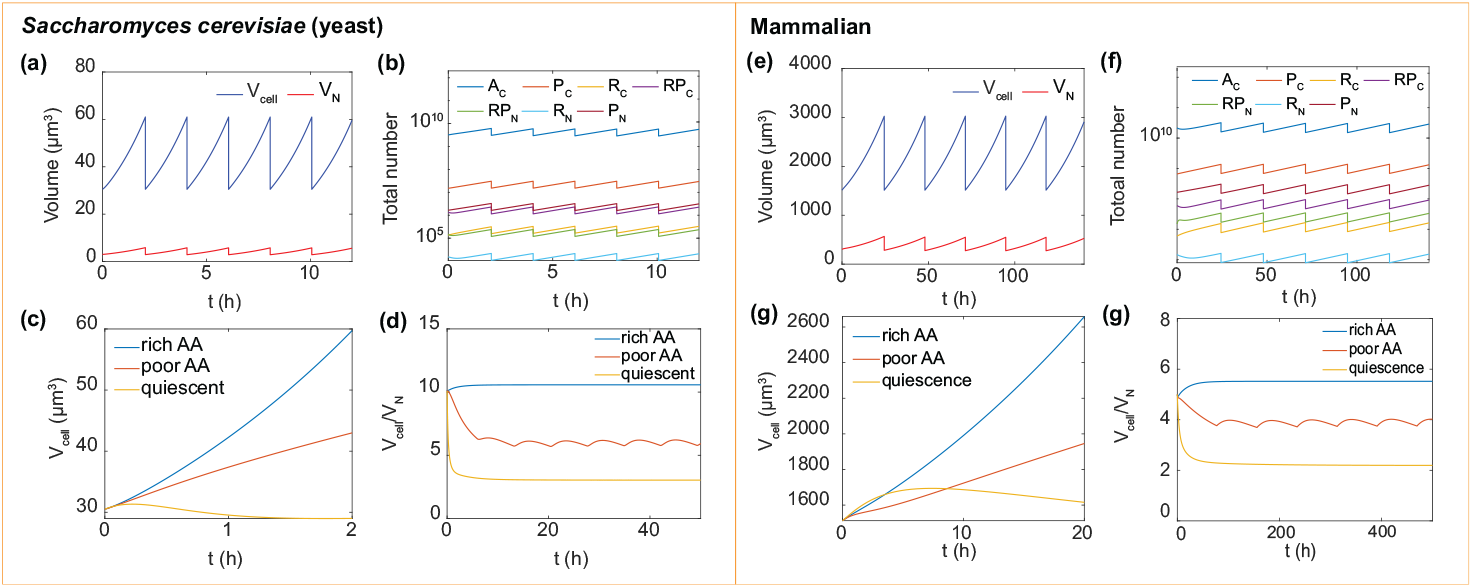
Model results for cell and nuclear volumes of *Saccharomyces cerevisiae* (yeast) and HeLa (mammalian) over several cell cycles. Here the cell divides when the cell volume doubles. (a) The cell volume and nuclear volume for a repeatedly dividing cell. (b) The number of cell components (amino acids, proteins, ribosomal proteins and ribosomes) for successive rounds of cell division. (c) The cell volume grows exponentially if amino acid is rich, and 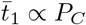. The growth is linear if amino acid is poor and 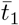 is a constant. In the quiescent case, the cell volume becomes stationary. (d) In all cases, the C/N ratio reaches a constant. In the quiescent case, the cell growth rate is zero, and the C/N ratio is small. (e)-(h) Corresponding results for mammalian cell.

Fig. 4(c,g) show cell growth trajectories in rich amino acid, poor amino acid and non-growing (quiescent) conditions. In the nutrient-limited case, the transport rate is set to be a constant, 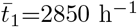 for the yeast cell and 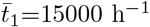 for the mammalian cell. In the stationary case, the cell is usually in an amino-acid-limited environment, with significantly decreased protein synthesis and increased degradation [35, 36]. In this case, we assume that proteasomes and lysosomes are sufficient and the degradation (disassembly) rate is only proportional to the number of proteins or ribosomes to be degraded (disassembled) (i.e., −*dP/dt* = − *d* ×*P, dR/dt* = *d* ×*R*). All the degradation coefficients are set to be *d* = 0.8 for yeast cell and *d* = 0.05 for mammalian cell. For yeast cell, the transport rate is 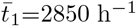 and ribosome synthesis rate is set to be *s*_2_ = 5 (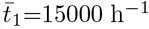 and *s*_2_ = 0.3 for mammalian cell). Compared with the cell in rich-nutrient environment (exponential growth), in poor-nutrient environment, our model predicts that the cell grows linearly due to limited supply of amino acids. In the stationary case, the cell almost maintains a constant volume due to the balance between synthesis and degradation.

### Cytoplasm to nucleoplasm volume ratio

Consistent with observations, our model predicts a constant cytoplasm to nucleoplasm volume (C/N) ratio for all cases, but the value of the ratio varies significantly for different cells in different conditions. Fig. 4(d,g) show the change of cell to nucleus volume ratio (C/N ratio) in several cell cycles in the three cases. For both rich-amino acid and quiescent cases, C/N ratios reach a constant (10 and 3 for yeast cell; 5.5 and for mammalian cell). The C/N ratio is low in quiescent cell because there are relatively more proteins in nucleus due to nuclear import and decreased ribosome synthesis. For the nutrient-limited cell, the C/N ratio fluctuates periodically. This is because in one cell cycle, the linearly growing cell is unable to adjust all the components into a stable state. At the beginning of the cycle, C/N ratio increases, then it decreases because imported amino acids are not sufficient to maintain increased cytoplasmic protein synthesis. This result comes from the assumption that amino acid import is constant throughout the cell cycle, but it is also possible that the cell adjusts amino acid import as the cell cycle progresses. After each division, the cell state is reset. Notably, in the three cases above, protein distributions in cytoplasm and nucleoplasm are totally different due to different synthesis and transport, and this is a major reason for the different C/N ratios in these cases.

### Effects of protein transport and synthesis on cell growth rate

Fig. 5(a)-(e), (k)-(l) show how model parameters influence the cell growth rate, *λ*, in exponentially growing cells (rich-amino acid case) for both yeast and mammalian cells. The amino acid import coefficient *t*_1_ increases the growth rate (Fig. 5(a,k)). When *t*_1_ is low, the growth rate can be negative because there is more protein degradation than synthesis. Ribosomal protein and ribosome transport coefficients *t*_2_ and *t*_3_ both increase the growth rate when they are small. However, when they are large, *t*_3_ has little influence, while *t*_2_ decreases the growth rate due to a significant decrease of cytoplasmic ribosomal proteins (Fig. 5(b,l)). Compared with other transport activities, transport of non-ribosomal proteins into (*t*_4_) and out of (*t*_5_) nucleus is more important. There is a sharp transition around the line *t*_4_ = *t*_5_. when *t*_5_ is greater than *t*_4_, the growth rate keeps constant; otherwise, the growth rate declines significantly to zero (Fig. 5(c,m)). This sharp change is due to the sharp change of transporter proteins. Increase of *t*_4_ significantly decreases the cytoplasmic protein amount, thus decreasing the transport-related proteins in the cytoplasm proportionally. As a result, the ribosome synthesis cycle is slowed down, thus limiting the growth rate. On the other hand, with increasing *t*_5_, the transported proteins accumulate. However, the growth rate is already “saturated” with the ribosomal protein and ribosome transports, so accelerating the ribosome cycle influences the growth rate very little. This result implies that the cell has to accurately control protein transport across the nuclear envelope.

**Fig 5.**
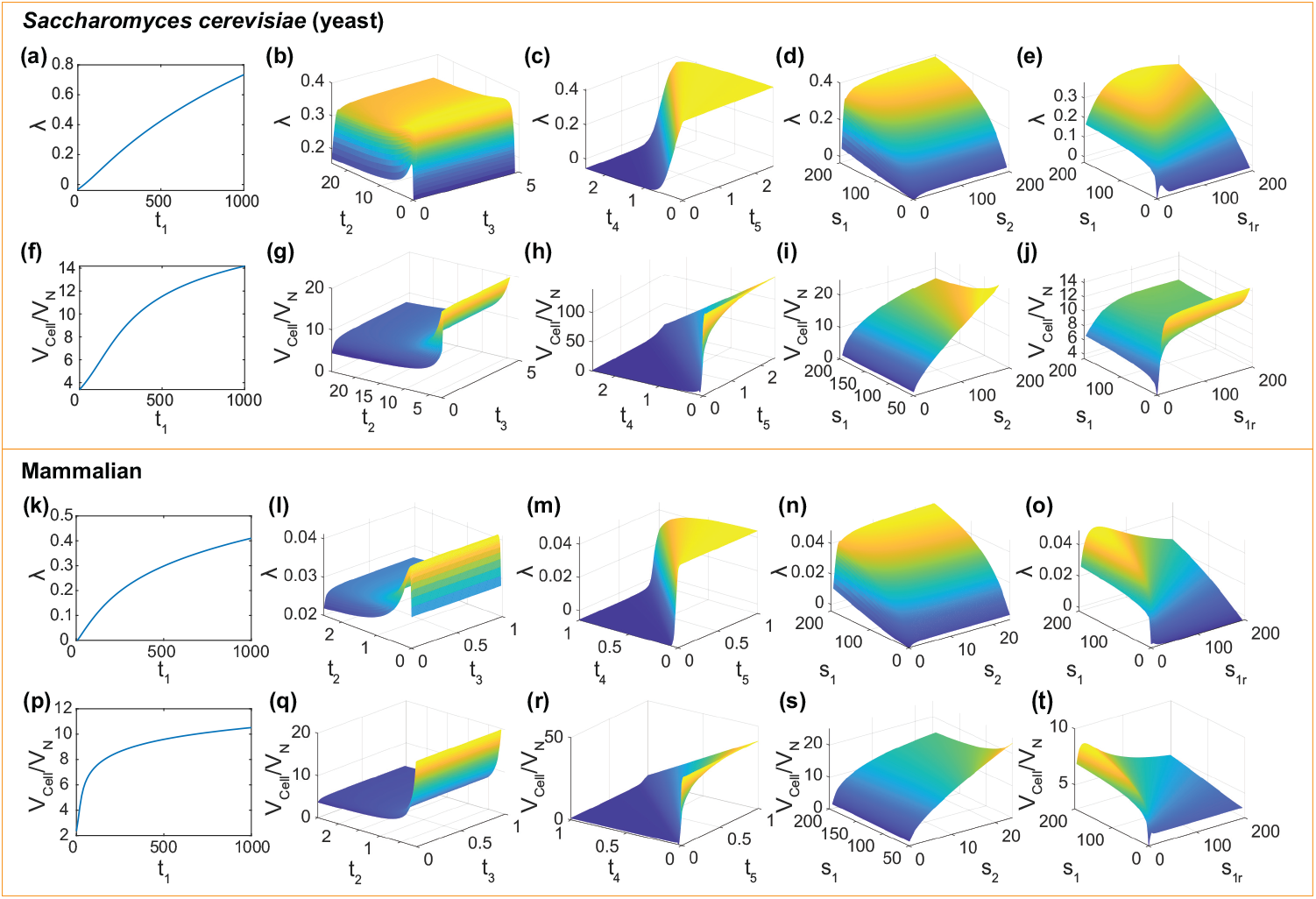
Effects of synthesis and transport parameters on the cell growth rate and the C/N ratio. (a)-(c) Effects of amino acid import (*t*_1_), ribosomal protein transport (*t*_2_), ribosome transport (*t*_3_) and non-ribosomal protein transport (*t*_4_ and *t*_5_) on growth rate. Notably, the growth rate is non-monotonically influenced by *t*_2_, and there’s a sharp change in growth rate around *t*_4_ = *t*_5_. (d)-(e) Effects of non-ribosomal and ribosomal protein synthesis (*s*_1_ and *s*_1*r*_) and ribosome synthesis (*s*_2_) on growth rate. There is an optimal *s*_1_*/s*_1*r*_ for the maximum growth rate. (f)-(j) Effects of transport and synthesis on the C/N ratio. protein synthesis coefficients *s*_1_ and *s*_1*r*_ have a non-monotonic influence on C/N ratio. (k)-(t) Corresponding results for mammalian cells. Interestingly, compared to yeast cell case, *s*_1_ and *s*_1*r*_ have different effects on C/N ratio due to different ribosome synthesis coefficient *s*_2_.

Non-ribosomal protein synthesis coefficient *s*_1_ and ribosome assembly coefficient *s*_2_ both increase the growth rate. However, *s*_1_ has a more obvious influence than *s*_2_ (Fig. 5(d,n)). The ribosomal protein synthesis coefficient *s*_1*r*_ also plays an important role in cell growth. Both synthesis coefficients *s*_1_ and *s*_1*r*_ increase the growth rate. However, there is also a trade-off between *s*_1_ and *s*_1*r*_, which is indicated by the non-monotonicity of growth rate when fixing *s*_1_ + *s*_1*r*_ (Fig. 5(e,o)). Therefore, there is an optimal ribosomal and non-ribosomal protein synthesis ratio for the maximum growth rate. This non-monotonicity also agrees with findings in bacteria [37].

### Effects of protein transport and synthesis on C/N ratio

Fig. 5 (f)-(j), (p)-(t) show how model parameters influence the C/N ratio for both yeast and mammalian cells. In exponentially growing cells, amino acid uptake coefficient *t*_1_ increases the C/N ratio because the amino acid import rate is much more than the protein transport rate across nuclear membrane, thus causing the accumulation in the cytoplasm (Fig. 5(f,p)). The ribosome transport coefficient *t*_3_ increases the C/N ratio when it is small, but has little influence when large (Fig. 5(g,q)). In contrast, the ribosomal protein transport coefficient *t*_2_ decreases the C/N ratio and it is more significant (Fig. 5(g,q)). Similar to growth rate, the non-ribosomal protein transport coefficients across nuclear membrane (into: *t*_4_, out of: *t*_5_) also influence the C/N ratio significantly, especially around *t*_4_ = *t*_5_ (Fig. 5(h,r)).

Interestingly, in addition to transport, synthesis also influences the C/N ratio. When the ribosome assembly rate coefficient *s*_2_ is high, the ribosomal proteins are rapidly converted to ribosomes, causing a rapid decrease in macromolecule number in the nucleus, thus resulting in a small nuclear volume and high C/N ratio (Fig. 5(i,s)). For yeast and mammalian cells, non-ribosomal protein synthesis coefficient *s*_1_ and ribosomal protein synthesis coefficient *s*_1*r*_ have different effects on the C/N ratio due to different *s*_2_ (Fig. 5(j,t)). For yeast cells, *s*_2_ is large, and ribosomal proteins in the nucleus are assembled at a high rate. Therefore, nuclear ribosomal proteins only take up a small portion of the nuclear volume. When increasing *s*_1*r*_, *RP*_*C*_ increases much more than *RP*_*N*_, so *s*_1*r*_ increases the C/N ratio. For a realistic ribosomal synthesis rate (*s*_1*r*_ = 125), *s*_1_ decreases C/N ratio. When *s*_1_ is small, ribosomal proteins dominate the cell volume. Due to large *s*_2_, the C/N ratio is high. However, when *s*_1_ is large, non-ribosomal proteins take a major part, and the influence of *s*_2_ on the C/N ratio is small. Non-ribosomal proteins are rapidly imported into the nucleus and cause a decrease in the C/N ratio. However, for mammalian cells, *s*_2_ is much smaller than for yeast, therefore *s*_1_ and *s*_1*r*_ have opposite effects on the C/N ratio.

### Effects of degradation and disassembly

We first consider the influence of protein degradation on the C/N ratio and the cell growth rate. Non-ribosomal protein degradation (*d*_1_) monotonically decreases growth rate and increases C/N ratio for both yeast and mammalian cells (Fig. 6 (a,e)).

**Fig 6.**
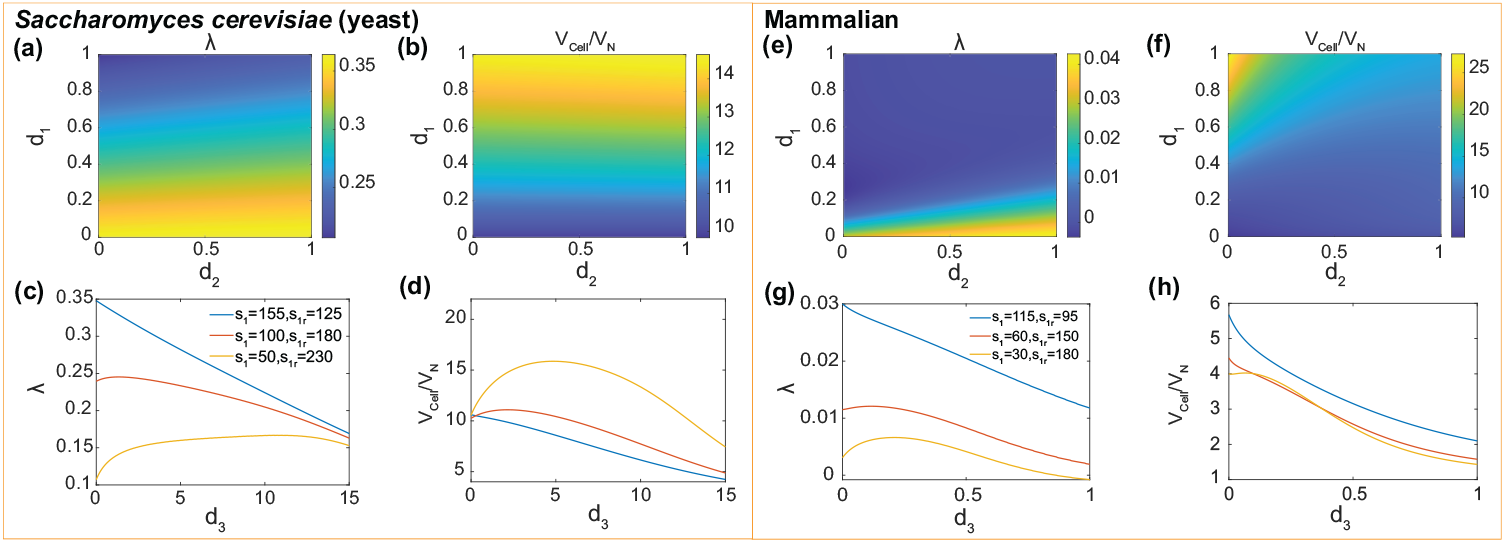
Effects of protein degradation and ribosome disassembly on cell growth and the C/N ratio. (a)-(b) effects of degradation on growth rate and C/N ratio. (c)-(d) Non-monotonic influences of disassembly on growth rate in different ribosomal protein synthesis levels. (e)-(h) Corresponding results for mammalian cells.

Ribosomal protein degradation (*d*_2_) has little influence on the growth rate. However, *d*_2_ decreases the C/N ratio for mammalian cell when *d*_1_ is large (Fig. 6 (b,f)).

In contrast from protein degradation, ribosome disassembly has two effects on cell growth: on the one hand, the disassembly of ribosome produces more ribosomal proteins (*R*_*C*_ → *n*_3_*RP*_*C*_), thus increasing the cell volume; on the other hand, disassembly decreases the protein synthesis rate. With competition between these two effects, we see that as the ribosomal and non-ribosomal protein ratio varies, disassembly has different influences on cell growth (For simplicity, we assume the degradation rates are the same in cytoplasm and nucleus. i.e., *d*_3*C*_ = *d*_3*N*_ = *d*_3_). For both yeast and mammalian cells, for high non-ribosomal protein ratio: *s*_1_ = 155, *s*_1*r*_ = 125 for yeast cell and *s*_1_ = 115, *s*_1*r*_ = 95 for mammalian cell, *d*_3_ decreases the growth rate monotonically. However, when *s*_1*r*_*/s*_1_ is increased, *d*_3_ changes the growth rate non-monotonically (Fig. 6(c,g)). When *d*_3_ is low, it increases the growth rate; when it is high, it decreases the growth rate. These results can be explained as follows: When *s*_1_*/s*_1*r*_ is high, non-ribosomal protein dominates the cell volume. With increase of ribosome disassembly, ribosome number decreases, thus decreasing the non-ribosomal protein synthesis rate, and this influence is larger than the volume increase caused by *RP* increase. However, when *s*_1_*/s*_1*r*_ is low, the ribosomal protein amount increases and also becomes important for determining cell volume, and the increase of *RP* number influences more than the growth decrease caused by less ribosomes. It is worth noting that when *d*_3_ is too high, it also decreases the ribosomal protein amount due to insufficient ribosomes. As a result, *d*_3_ finally decreases the growth rate. Ribosome disassembly has similar influence on the C/N ratio (Fig. 6(d,h)).

### Effects of ribosomal protein to ribosome ratio

It is suggested that in aneuploid cells, cell volume abnormality might be related to the abnormal ratio of protein complex amount to free protein amount [38]. The ribosome is a major protein complex in the cell, and the ratio of ribosome to ribosomal protein might be an indicator of aneuploidy. Therefore, it is necessary to explore how the ratio RP/R influences cell growth. In our model, there are several ways to tune this ratio: Generally, *d*_3_ increases the ratio, while *s*_2_ or *t*_2_ decreases this ratio. When tuning *d*_3_, the effect of RP/R on growth rate is similar to that of *d*_3_ (Fig. 7(a,d)). In the process of tuning *s*_2_, when RP/R is low (*s*_2_ is high), the growth rate decreases with this ratio. At this state, non-ribosomal proteins dominate the cell volume, and the increase of RP/R decreases the non-ribosomal protein synthesis by decreasing the ribosome number, thus decreasing the growth rate. However, when RP/R is high enough (*s*_2_ is low), ribosomal proteins play a more important role in cell volume, so RP/R slightly increases growth rate by increasing the ribosomal protein number (Fig. 7(b,e)). When tuning *t*_2_, RP/R initially increases and then decreases the growth rate (Fig. 7(c,f)). Generally speaking, the non-monotonic change of growth rate is determined by the trade-off between ribosomal protein increase (increasing macromolecule number, thus cell volume) and cytoplasmic ribosome decrease (decreasing protein synthesis). In different situations, the ribosomal proteins and cytoplasmic ribosomes play different roles, thus the RP/R-*λ* curve have nontrivial shapes.

**Fig 7.**
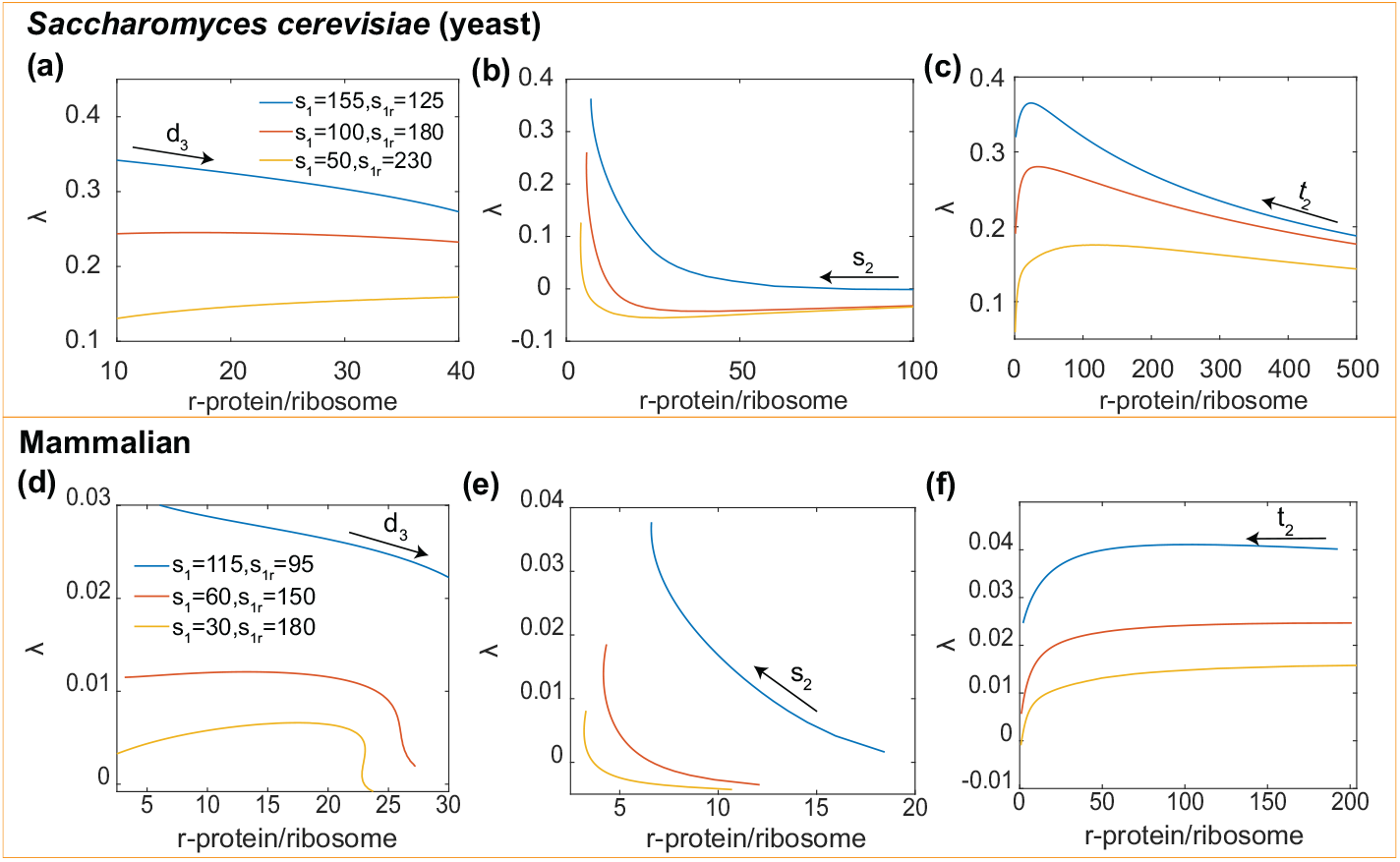
Non-monotonic effects of ribosomal protein to ribosome ratio (RP/R) on growth rate when tuning (a) disassembly coefficient *d*_3_, (b) ribosome synthesis coefficient *s*_2_ and (c) ribosomal protein synthesis coefficient *t*_2_. (d)-(f) Corresponding results for mammalian cells.

### Optimal strategies for regulating cell growth and C/N ratio

We have seen that even for this simple model, many parameters can influence the cell growth rate and the C/N ratio, it is natural to ask whether the cell controls these parameters in groups, for example using canonical growth pathways. If so, does the cell have different strategies to tune the parameters under different situations? To answer these questions, we firstly normalize the parameters by their realistic values (e.g., 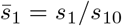), and then calculate the gradients of the cell growth rate and the C/N ratio with respect to the normalized model parameters: e.g., 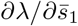. The gradient vector is then normalized by its length. This can be done for the exponential growth (rich AA) and linear growth (poor AA) cases. For the quiescent case, we compute the gradient of the steady state volume with respect to parameters. Results are presented in radar charts, and the +/-sign means positive/negative derivatives (Fig. 8), i.e., whether the parameter has a positive or negative influence on the quantity considered.

**Fig 8.**
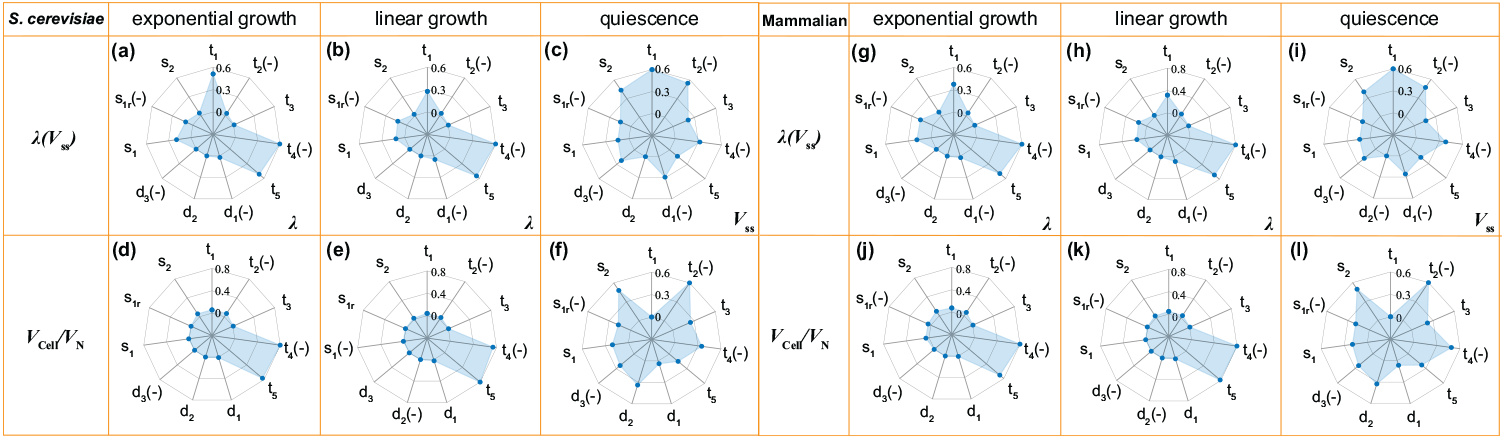
(a)-(f) Gradients of the cell growth rate and the C/N ratio with respect to the normalized model parameters for exponential, linear growth conditions, respectively. For the quiescent case, gradients of the steady state cell volume with respect to parameters are shown. Amino acid and non-ribosomal protein transport (*t*_1_, *t*_4_, *t*_5_) are most important parameters in exponential and linear growth conditions. For the quiescent case, ribosomal protein transport (*t*_2_) and ribosome synthesis (*s*_2_) play most important roles. (g)-(l) Corresponding results for mammalian cell.

For both mammalian and yeast cells, when extracellular amino acid is saturating, the cell grows exponentially, and the growth rate is influenced mostly by non-ribosomal protein transport coefficients *t*_4_, *t*_5_ and amino acid import coefficient *t*_1_ (Fig. 8(a,g)), so tuning nutrient uptake rate and reallocating protein distribution in cytoplasm and nucleus is the most effective way to adjust growth rate. For the C/N ratio, *t*_4_, *t*_5_ are the most influential parameters (Fig. 8(d,j)). The AA-limited case is similar to the rich AA case, but amino acid import (*t*_1_) plays a less important role in tuning cell growth (Fig. 8(b,e,h,k)). In the quiescent case, the cell enters a steady state with a constant cell volume. Compared with the other two cases, ribosomal protein import (*t*_2_) and ribosome synthesis (*s*_2_) become the speed limiting steps in the ribosome cycle.

Therefore, besides amino acid uptake (*t*_1_), *t*_2_ and *s*_2_ are the most important parameters influencing the steady state volume and C/N ratio (Fig. 8(c,f,i,l)).

In summary, in three growth conditions considered, by balancing resource uptake, allocation and utilization, the cell requires different strategies to control growth rate (steady state volume) and C/N ratio. For growing cells, besides amino acid import *t*_1_, non-ribosomal protein transport coefficients *t*_4_ and *t*_5_ are also two of the most important parameters. These two parameters significantly influence the cytoplasmic transport protein amount, which further influence the speed of the ribosome assembly cycle. In such a case, growth resources are rich, and the determining factor of the growth is the allocation of these resources in cytoplasm and nucleus. However, in quiescent case, due to low ribosome synthesis, ribosomal protein import (*t*_2_) and ribosome synthesis (*s*_2_) become the speed limiting steps and have more effects in the ribosome assembly cycle and cell steady state.

### Distributions of cell volume and proteome in a growing population

With the advent of single-cell techniques, it is becoming possible to analyze the single cell volume and the single cell proteome composition. The data can be assembled to obtain a cell volume or proteome distribution for a particular cell type. Our model can be used as a starting point to obtain cell volume and proteome distributions.

Mathematically, Eqs. (23-29) are velocities of protein level and cell size increase. These velocities can be used in a stochastic model to generate cell volume and proteome distributions which has been discussed elsewhere [39, 40]. For example, a computation of protein distribution for each cell goes as follows (Fig. 9(a)): (1) start from *N* initial cells with model variables evenly distributed between 1 and twice the estimated initial conditions; (2) At each time step, increment the model variables (*A*_*C*_, *P*_*C*_, *RP*_*C*_, *R*_*C*_), etc, according to governing velocities in Eqs. (23-29) for each time step *dt* = 0.02. Then calculate the division probability density as a function of cell volume *V*_*C*_ as [39]:

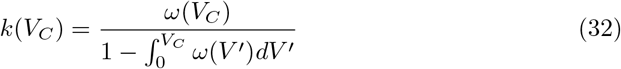

where *ω*(*V*_*C*_) is the division volume distribution; (3) Determine whether the cell divides during this time step: if 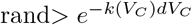, the cell divides symmetrically with some noise, and the volume difference between newborn cells follows a normal distribution: Δ*V*_*new*_*/V*_*div*_ ∼ *N* (0, 0.125) (all the constituents are divided following the same distribution) [41]; otherwise, the variables increase after the time increment. All the parameters in the model are the same as other sections. After a number of iterations, the volume and protein number distributions equilibrate to a steady distribution.

**Fig 9.**
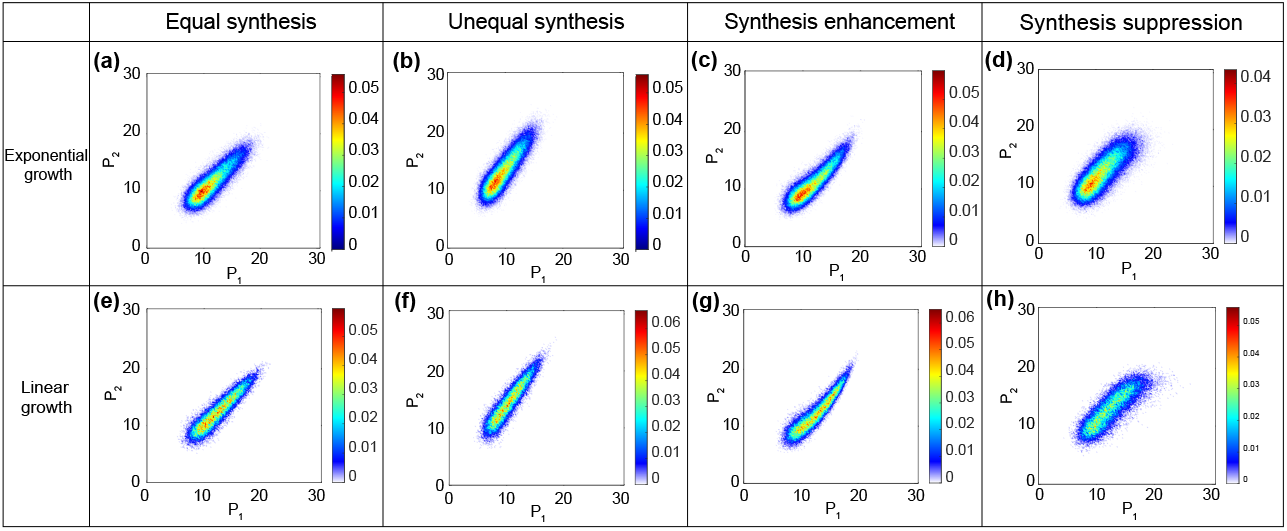
Computed proteome distributions (yeast parameters) in exponential growth and linear growth. Mammalian cell models show a similar behavior. (a) equal synthesis of *P*_1_ and *P*_2_ without gene regulation. The two proteins are naturally correlated because of growth through cell cycle. (b) unequal synthesis without correlation. (c) one protein enhances the synthesis of the other. (d) one protein suppresses the synthesis of the other. The distribution is not linearly correlated for synthesis enhancement and suppression cases. (e)-(h) Corresponding results for linear growth.

In a real cell, different proteins are translated at different rates. *s*_1_ depends on the mRNA level of a particular protein, and gene regulatory interactions that may enhance or suppress the translation this particular protein. Indeed, *s*_1_ should be proportional to the mRNA concentration, which can be measured in single cell RNAseq [42]. However, even without gene regulation and as long as the protein is actively transcribed throughout the cell cycle, we see that the protein distributions are naturally correlated, even when there are no gene regulatory interactions (Fig. 9 (a,e)).

To study how enhancement or suppression of expression can influence the proteome distribution, we include 2 types of non-ribosomal proteins in our model, where one protein influences the synthesis of the other. The synthesis rates are *s*_1,1_, *s*_1,2_, respectively. Further, we take yeast cell as an example and calculate the distributions under different synthesis situations in exponentially and linearly growing cells using the velocities from our model. In the first case, two proteins have the same synthesis rate and do not effect each other’s translation. In the second case, protein *P*_1_ and *P*_2_ have different synthesis rates (*s*_1,1_ = 65, *s*_1,2_ = 90), but do not effect each others translation. In the third case, *P*_1_ enhances the synthesis of *P*_2_: *s*_1,2_ = 6*P*_1_. In the fourth case: *P*_1_ suppresses the synthesis of *P*_2_: *s*_1,2_ = 145 − 5*P*_1_. The computed protein distributions are shown in (Fig. 9(a)-(d)) for exponential growth and (Fig. 9(e)-(h)) for linear growth.

Fig. 9(a),(b),(e),(f) show that even without gene regulatory interactions, the distribution of two proteins are naturally correlated. This is because there is an overall increase of all proteins during cell growth and cell cycle. For equal synthesis rates, the slope of the correlation is 1 while if the synthesis rate are unequal, the slope can be any value. With synthesis enhancement or suppression, we can see that the distributions are more skewed and are not linearly correlated (Fig. 9(c)-(d),(g)-(h)). Based on Eq. (2), by assuming constant ribosome concentration and equal growth rates for all the cell components, the correlation relationships between the two proteins can be approximated by: *P*_2_ ∝ *P*_1_^2^ and *P*_2_ ∝ *P*_1_ − *P*_1_^2^ for synthesis enhancement and suppression, respectively.

Note that these computed proteome distributions will depend on the division mechanism. We have used a division probability based on Eq. 32. Other mechanisms will in general give different distributions. In principle, our model can be refined to include all protein species and examine division mechanisms based on the proteome composition and cell volume.

## Discussion and Conclusions

In this work, we developed a coarse-grained eukaryotic cell growth model, which includes protein and ribosome synthesis, transport across both cell surface and the nuclear envelop, protein degradation and ribosome disassembly. We simulate cell growth for yeast and mammalian cells in three different amino acid conditions: rich-AA, poor-AA and the quiescent case. When amino acids are abundant, cells grow exponentially. However, when AA is poor, cells grow linearly due to limited AA import. When the cell enters the quiescent state, it gradually stops growing and cytoplasmic proteins are rapidly transported into nucleus, giving a low C/N ratio. In the rich-AA and quiescent cases, the C/N ratio reaches constant, but in the poor-AA case, the C/N ratio can fluctuate. Furthermore, we studied how transport, synthesis and degradation parameters influence the growth rate and the C/N ratio. Growth rate is dependent on transport across nucleus membrane. In particular, growth rate is found to be highly sensitive to non-ribosomal protein transport (*t*_4_ and *t*_5_), which means the cell should have tight control over these parameters. For a given total protein production rate, there exists an optimal ratio between ribosomal and non-ribosomal protein synthesis, at which the cell growth rate can achieve maximum, similar to what has been found in bacterial cells [37]. We also show that in addition to transport, protein synthesis parameters also influence the C/N ratio significantly. Ribosome disassembly coefficient *d*_3_ can influence both the growth rate and the C/N ratio non-monotonically, which is due to the trade-off between disassembly-caused macromolecule number increase and cytoplasmic ribosome number decrease. Moreover, we show that the ribosomal protein to ribosome ratio RP/R also influences the growth rate non-monotonically, which could explain the size abnormality and growth rate abnormality in aneuploid cells.

It should be noted that all the kinetic parameters in the model are likely to be actively controlled in cells. Canonical growth pathways such as the AKT [43], TGF-beta [44], MTor [45] and Hippo pathways [46], and nuclear transport pathways such as the RanGTP cycle have been identified to influence cell growth and nucleoplasm-cytoplasm transport. The influence of these pathways on model parameters have yet to be studied systematically. Moreover, in different growth environments, our model predicts that the cell should utilize different strategies to achieve different growth rate or steady state volume. For exponentially or linearly growing cells, amino acid and non-ribosomal protein transport are the most important parameters, while amino acid import coefficient *t*_1_ is more important for exponentially growing cells. However, for quiescent cells, ribosomal protein import and ribosome synthesis coefficients become the most important parameters. These strategies for optimal growth apply for both yeast and mammalian cells. The identified growth pathways are likely to utilize feedback mechanisms to control cell growth. This opens the possibilities of non-monotonic changes in the growth rate: for instance, time delays in the feedback mechanisms will introduce growth rate oscillations [47].

Experiments also shows considerable randomness in the cell growth rate, the cell volume and the C/N ratio, even for isogenic cells at the same point in the cell cycle. This suggests that there is significant stochasticity in all the parameters in the model. The significances of this stochasticity is unclear, but could be related to how stochastic dynamical systems response to changes in environmental conditions. A system exhibiting more randomness also responses to changes faster [48]. This allows the dynamical system to rapidly adapt to new environments. Therefore, the observed cell size and growth fluctuations may be a feature of cell control system that regulate the cell physiological state.

Our model can be generalized to examine specific proteins with differing synthesis rates. The results can be compared with single cell proteomic data. By examining a model with 2 proteins, we find that the computed proteome distribution will have a trivial correlation if the proteins are synthesized independently. This is because of cell cycle effects where older cells will generally have larger number of proteins. However, if the the proteins can mutually enhance or suppress their expression, additional correlations will show up in the proteome distribution. From single cell proteomics data, our model maybe used to infer gene regulatory interactions.

An interesting aspect of the model is that it predicts cell mechanical behavior can potentially couple to cell growth. There are several places where this potential coupling can occur. One, from studies of cell water and ion homeostasis, the cell volume is predicted to be proportional to the total number of macromolecules in the cell (proteins and protein complexes). The proportionality constant, however, depends on the cytoplasmic hydraulic pressure and therefore could depend on cytoskeletal contractility. Two, water and ions are generally thought to freely diffuse across the nuclear pores. However, because the nuclear envelope is mechanically connected to the cytoskeleton through LINC complexes, there may be additional mechanical forces on the nuclear envelope that balances increased hydraulic and osmotic pressures in the nucleus. Moreover, mechanical forces on the nuclear envelope may also alter nucleoplasm-cytoplasm transport rate by changing the tension in the nuclear membrane and the nuclear pore complex. Finally, direct connection between cell surface tension and cell growth pathways has been found for yeast and mammalian cells. Therefore, cell mechanical behavior can directly influence cell growth. The coupling of growth to mechanics of the cell and tissue is fundamental for our understanding of morphogenesis and this paper outlines a framework where force-dependent growth can be examined in more detail.

## Acknowledgments

We would like to gratefully acknowledge Adrian Pegarano and David Weitz for providing the data for Figure 1. YFW and SXS are supported by NIH R01GM134542. PJ is supported by NIH R35GM136259.

